# Multiple known and a novel parvovirus associated with an outbreak of feline diarrhea and vomiting

**DOI:** 10.1101/2020.03.24.005876

**Authors:** Yanpeng Li, Emilia Gordon, Amanda Idle, Eda Altan, M. Alexis Seguin, Marko Estrada, Xutao Deng, Eric Delwart

**Author notes:** Correspondence: Eric Delwart, Vitalant Research Institute, 270 Masonic Avenue, San Francisco, CA 94118, United States. These authors contribute equally to the study (Yanpeng Li, Emilia Gordon).

## Abstract

An unexplained outbreak of feline diarrhea and vomiting, negative for common enteric viral and bacterial pathogens, was subjected to viral metagenomics and PCR. We characterized from fecal samples the genome of a novel chapparvovirus we named fechavirus that was shed by 8/17 affected cats and different feline bocaviruses shed by 9/17 cats. Also detected were nucleic acids from attenuated vaccine viruses, members of the normal feline virome, viruses found in only one or two cases, and viruses likely derived from ingested food products. Epidemiological investigation of disease signs, time of onset, and transfers of affected cats between three facilities support a possible role for this new chapparvovirus in a highly contagious feline diarrhea and vomiting disease.

## 1. Introduction

Cats have an estimated world-wide population of over half a billion. Members of at least 15 viral families have been found to infect cats, including rabies virus, feline rotavirus (FRV), feline protoparvovirus virus (FPV), feline bocaviruses (FBoV), feline astroviruses (FeAstV), feline picornaviruses (FePV), feline enteric coronavirus (FECV), feline calicivirus (FCV), feline herpesvirus (FHV-1), feline immunodeficiency virus (FIV) and feline leukemia virus (FeLV) (Di Martino, Di Profio, Melegari, & Marsilio, 2019; Ng et al., 2014; Pesavento & Murphy, 2014). Diarrhea in cats is common and possible infectious causes include bacteria, parasites, and/or viruses. Some of the most prevalent feline enteric viruses include FBoV, FeAstV, FRV, and FPV (Ng et al., 2014; Otto et al., 2015; W. Zhang et al., 2014). Conditions in animal shelters contribute to pathogen emergence due to factors including intensive housing, rapid population turnover, animal stress, and the presence of many direct and indirect routes of possible exposure (Pesavento & Murphy, 2014).

The *Parvoviridae* family consists of non-enveloped, icosahedral, viruses with single stranded DNA genomes of 4 to 6Kb (Cotmore & Tattersall, 2014; Mietzsch, Penzes, & Agbandje-McKenna, 2019). Eight ICTV approved genera are currently included in the *Parvoviridae* family (Cotmore et al., 2019). A recent reorganization of the *Parvoviridae* has proposed the creation of a third subfamily named *Hamaparvovirinae* that includes invertebrate and vertebrate viruses as well as endogenized genomes in the germ lines of fish and invertebrates suggesting an ancient origin (Penzes, de Souza, Agbandje-McKenna, & Gifford, 2019; Souza et al., 2017). Vertebrate infecting members of this subfamily are classified within the *Chaphamaparvovirus* genus (previously labeled as unclassified or chapparvoviruses) have been identified in rats and mice (Williams et al., 2018; Yang et al., 2016), bats (Baker et al., 2013; Yinda et al., 2018), rhesus macaques (Kapusinszky, Ardeshir, Mulvaney, Deng, & Delwart, 2017), cynomologus macaques (Sawaswong et al., 2019), dogs (Fahsbender, Altan, Seguin, et al., 2019), pigs (Palinski, Mitra, & Hause, 2016), Tasmanian devils (Chong et al., 2019)), birds (red-crowned crane (Wang et al., 2019), chicken (Lima et al., 2019), and turkey (Reuter, Boros, Delwart, & Pankovics, 2014)), and fish (Gulf pipefish (Penzes et al., 2019) and Tilapia (Du et al., 2019)). A recent study showed that a murine chaphamaparvovirus named murine kidney parvovirus (MKPV) was the cause of nephropathy in laboratory mice (Roediger et al., 2018) and was widely distributed world-wide (Christopher J. Jolly, 2019).

Here, we analyzed a multi facility outbreak of diarrhea and vomiting in cats using a commercial feline diarrhea panel of PCR tests for known enteric pathogens, viral metagenomics, and PCRs. Multiple mammalian viruses of varied origins were detected. Diverse feline bocaviruses and a novel chaphamaparvovirus we named fechavirus were each shed by approximately half of the affected animals tested. The epidemiology of the outbreak points to a possible role for this newly characterized parvovirus in feline diarrhea pending larger studies.

## 2. Materials and Methods

### 2.1. Sample collection and pathogen screening

From November 2018 to January 2019 a multifacility outbreak of feline vomiting and diarrhea occurred in an animal shelter system in British Columbia, Canada. The case definition was: cats housed in three affected facilities with one or more episodes of vomiting or diarrhea (with no apparent other cause) and direct or indirect exposure to cats from the case, between November 8, 2018 and January 28, 2019. Indirect exposure was defined as exposure via contaminated fomites, personnel, or exposure to a contaminated environment. In total 43 cats met the case definition. An outbreak investigation was performed. Stool samples from 21 cats (15 individuals, two pooled litters of 3 co-housed kittens each) were collected and submitted for a variety of diagnostic tests. A subset of these were submitted to IDEXX Laboratories, Inc (Sacramento, CA, USA). A comprehensive cat diarrhea pathogen screening panel was used to test for potential enteric pathogens. Fifteen samples were collected within five days (the other 2 were within 10 days) after onset of vomiting or diarrhea with treatments generally initiated after sample collection. The IDEXX comprehensive feline diarrhea panel tests for Clostridium perfringens alpha toxin, Clostridium perfringens enterotoxin, Campylobacter coli and Campylobacter jejuni, Cryptosporidium spp., Giardia spp., Salmonella spp., Tritrichomonas foetus, Toxoplasma gondii, FPV, FECV, and FRV using PCR and RT-PCR. Four feline vomit samples were also collected from a room of cats in the third affected shelter between January 20 and 28, 2019. Other diagnostic testing on a subset of animals including fecal flotation [12 cats] (for parasites), fecal antigen testing (helminth [9 cats], Giardia [9 cats], and feline parvovirus [3 cats]) failed to yield a causative agent.

### 2.2. Viral Metagenomics analysis

One gram of feces from each of the five cats were pooled and vortexed in 2ml phosphate buffer saline (PBS) with zirconia beads. Viral particles were then enriched according to previously described methods by filtration through a 0.45-µm filter and digestion of the filtrate with a mixture of nuclease enzymes prior to nucleic acid extraction. Nucleic acids were then amplified using random RT-PCR (Li et al., 2015). Illumina library was generated using the transposon based Nextera XT Sample Preparation Kit (Illumina, CA, USA) and sequenced on the MiSeq platform (2 × 250 bases, dual barcoding). Duplicate and low-quality reads were removed and the Ensemble program was used to assemble contigs. Both contigs and singlets were then analyzed using the BLASTx (v.2.2.7) to all annotated viral proteins sequences available in RefSeq of GenBank. The short reads sequencing data is available at NCBI Sequence Read Archive (SRA) under the BioProject number PRJNA565775 (Biosample accession SAMN13526805-13526821, and SAMN13672796).

### 2.3. Genome assembly of novel chaphamaparvovirus

Geneious R11 program was used to align reads and contigs to reference viral genomes and generate partial genome sequence. The genome gaps from the initial assembly were then filled by PCR whose products were Sanger sequenced.

### 2.4. Diagnostic PCR and prevalence

DNA was extracted from each sample using the QIAamp MinElute Virus Spin Kit (Qiagen, Hilden, Germany), and a semi-nested PCR assays was used for the detection of viral nucleic acids in each sample. For fechavirus the first round PCR primers FechaF1 (5’-GGTGCGACGACGGAAGATAT-3’) and FechaR1 (5’-CAACACCACCATCTCCTGCT-3’) amplified a 332bp region. Second round of primers: FechaF2 (5’-GCTGCAGTTCAGGTAGCTCA-3’) and FechaR1 amplified a 310bp region. The PCR conditions for both rounds are as follows: 1x PCR Gold buffer I, 0.2mM dNTPs, 0.4µM of each forward and reverse primers, 1U of Amplitaq Gold DNA polymerase (Applied Biosystems, MA, USA) and 2µl DNA template in a final 25µl reaction. The PCR programs for both rounds: 95°C 5min, 35 cycles for 95°C 15s, 58°C 30s and 72°C 30s, followed by an extension at 72°C for 7min. PCR products were verified by gel electrophoresis and Sanger sequencing. The primers for feline bocavirus are as follows: FBoV-F 5’- AGAACCRCCRATCACARTCCACT-’3 and FBoV-R 5’-TGGCRACCGCYAGCATTT CA-’3, the PCR conditions are same as described before (Q. Zhang et al., 2019). The primers for FCV are: calici-F 5’- GCAAAGGTGGCGTCAAACAT -’3 and calici-R 5’- GCAAAGGTGGCGTCAAACAT -’3. The PCR programs: 95°C 5min, 40 cycles for 95°C 30s, 55°C 40s and 72°C 40s, followed by an extension at 72°C for 10min.

### 2.5. Phylogenetic analysis

The NS1 and VP1 protein sequences were aligned using Muscle program in MAGA 7.0, and the Neighbor-Joining phylogenetic trees for both NS1 and VP1 protein sequences were generated using bootstrap method under Jones-Taylor-Thornton (JTT) model.

## Results

### Epidemic and clinical data

A vomiting and diarrhea outbreak was identified across three animal shelters in British Columbia, Canada lasting from Nov. 2018 to Jan. 2019. A total of 43 cats met the case definition (see methods). 17 samples from 19 cats (including a pool of samples from 3 cats), plus four vomit samples, were available for further studies.

The outbreak was first identified on November 24, 2018 in Shelter 2 when 8/12 cats housed in the main adoption room became sick with vomiting and diarrhea. Diet, environmental, and toxic causes of the outbreak were ruled out clinically and an outbreak investigation initiated. The first cases in Shelter 2 (#614, #853) had arrived from Shelter 1 via the organization’s animal transfer program on November 15, 2018 and were sick during or shortly after transport. Upon further investigation, the originating shelter, Shelter 1, had 11 more sick cats. On November 20, Shelter 3 became involved in the outbreak when a cat (#160) transferred from Shelter 2 became sick, several days after arrival and before the outbreak had been identified. Ultimately, a total 13 cats were affected in Shelter 1 in November, 17 cats were affected in Shelter 2 (November – January), and 13 cats were affected in Shelter 3 (November-January) (Figure 1 and Supplementary Table 1). Nearly all transmission was indirect (Figure 2); because of this, it was not possible to definitively determine which animals had been exposed, except in specific rooms where housing was communal or exposure was known to be widespread prior to introduction of control measures. Attack rates for these rooms were 66.7% (Shelter 2) and 83.3% (Shelter 3) (Table 1).

**Table 1.** Attack rates (AR) for selected rooms and dates where widespread or complete exposure was suspected.

| Exposed (presumptive) |  |  |  |  |
| --- | --- | --- | --- | --- |
| Location | Sick | Not sick | Total | AR% |
| Shelter 2* | 8 | 4 | 12 | 66.7% |
| Shelter 3** | 10 | 2 | 12 | 83.3% |

\*November 23, 2018: Main adoption room
\*\* December 6, 2018-Jan 23, 2019: Communal cat area

**Figure 1.**
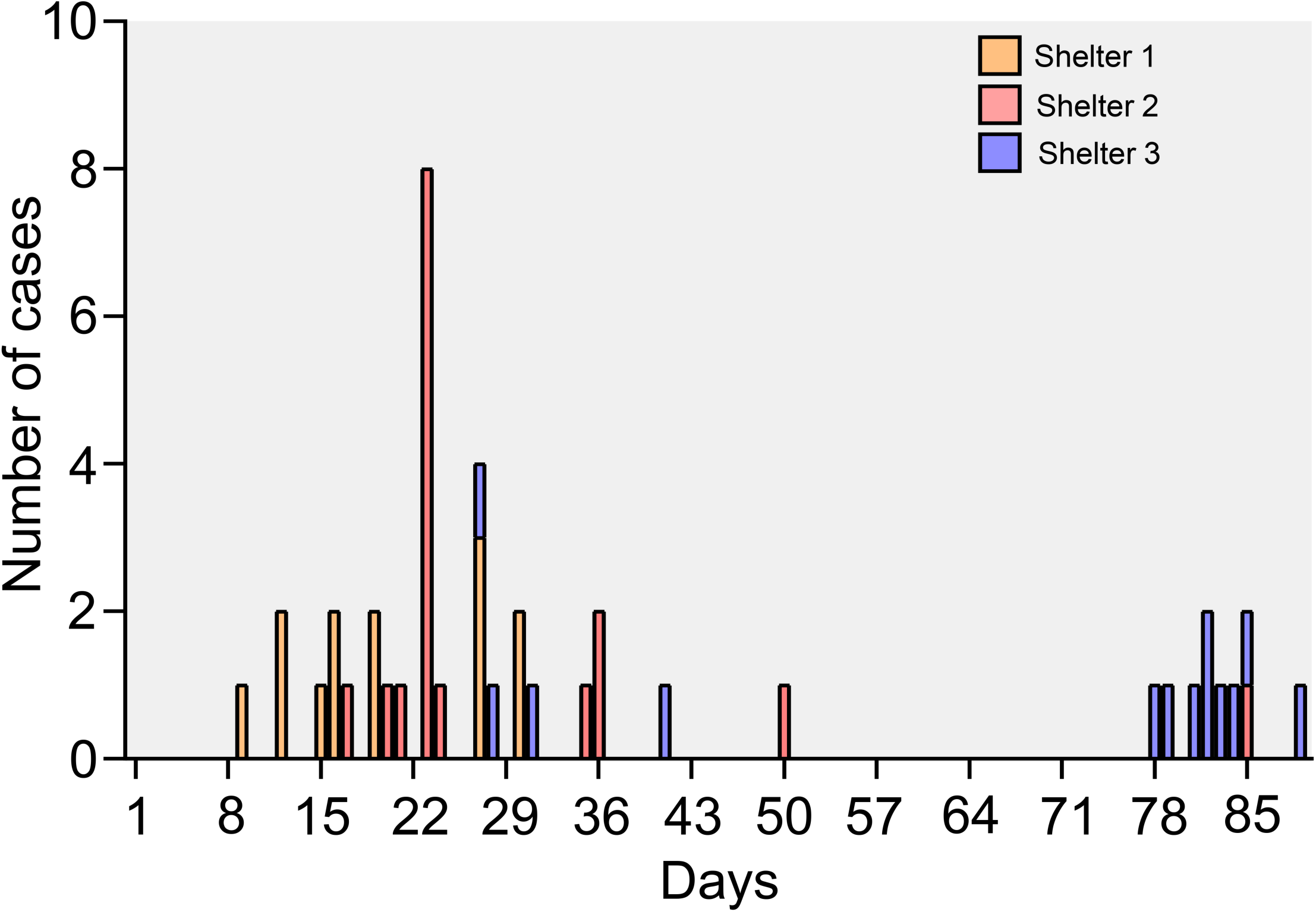
Daily epidemic cases of the cat diarrhea outbreak in three shelters from November to January.

**Figure 2.**
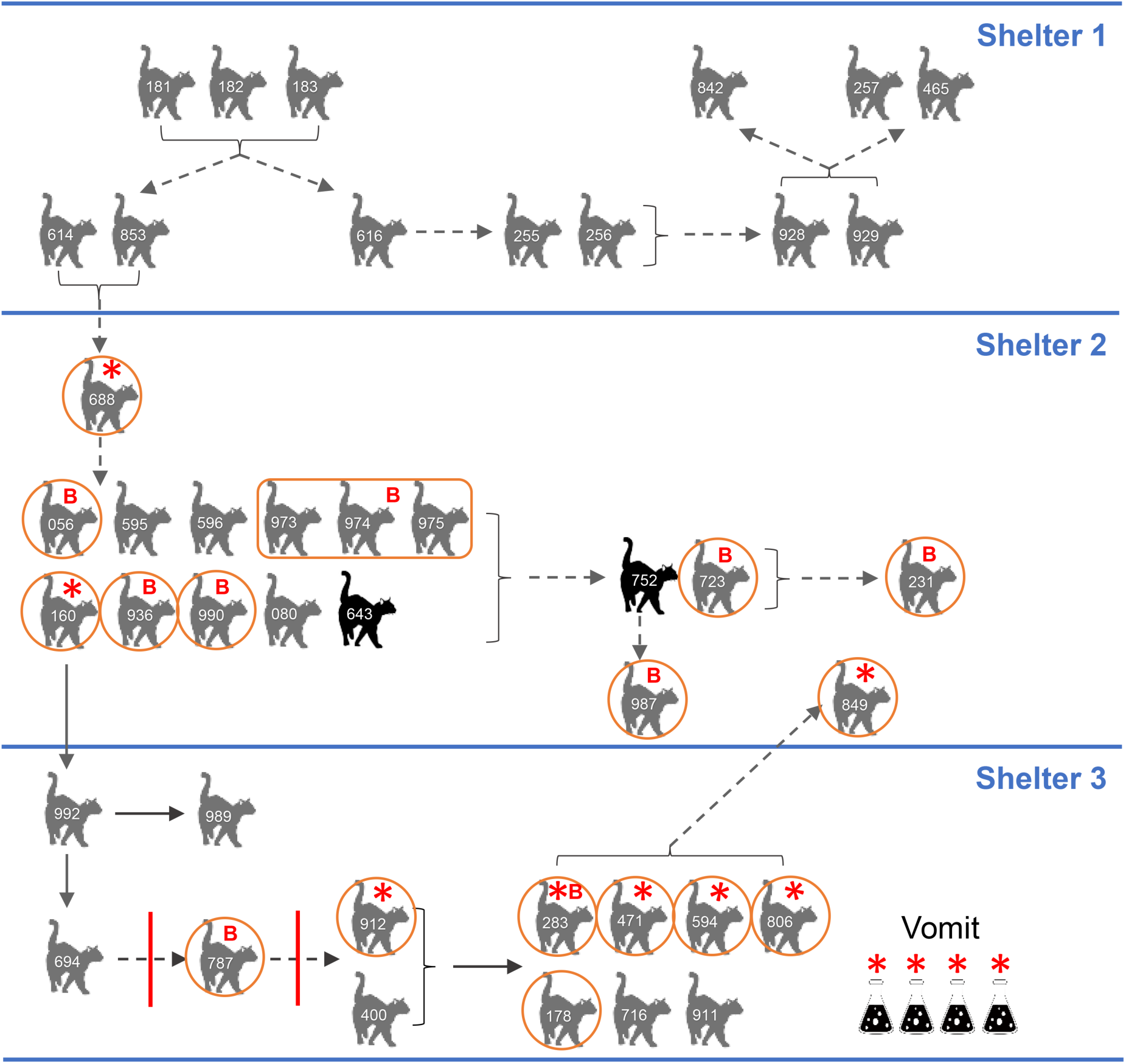
Outbreak diagram in three shelters. Animal ID shown for each cat (also see supplementary table). Grey cats: clinically affected and recovered, black cats died. Red circle or around each cat means sample was sampled individually, rectangle around the three cats means the cats were sampled together. Solid arrow means direct or indirect contact possible (housed communally), dotted line means indirect contact only. Bracket indicates group of affected animals with transmission to another group. Red solid vertical line indicates failed clean break. Red star means fechavirus PCR positive cats/samples. Red letter “B” means FeBoV PCR positive cats/samples.

Overall, diarrhea and vomiting were observed in 81.4% and 67.4% of the 43 cases. There were likely more cats vomiting because vomitus that could not be attributed to a particular cat was found multiple times in the final wave of illness. 25.6% and 11.6% of the affected cats also showed inappetence and lethargy, and 67.4% required veterinary care (Table 2). The minimum incubation period was 24 hours, and the maximum was estimated at 5-7 days based on estimated exposure dates. Vomiting tended to start 1-2 days before diarrhea, and last only a couple of days, but in some animals the diarrhea lasted up to a week (longer in a few animals). The mean duration of illness was 5.1 days and the median was 4.0 days (range 1 to 19 days) (Supplementary Table 1). No recovered cats relapsed and there were no cases where transmission was traced to a clinically recovered animal.

**Table 2.** Summary of clinical signs and need for outside veterinary care for 43 cats meeting case definition.

| Clinical signs | #of cats<br>(n=43) | Percentage |
| --- | --- | --- |
| Diarrhea | 35 | 81.4% |
| Vomiting | 29 | 67.4% |
| Inappetence | 11 | 25.6% |
| Lethargy | 5 | 11.6% |
| Required veterinary visit | 29 | 67.4% |

The sheltering organization initiated control measures on November 24, 2018 including cessation of all cat movement, personal protective equipment (gowns, gloves, caps, shoe covers) in all cat housing areas, and enhanced sanitation measures using accelerated hydrogen peroxide, which has good efficacy against bacteria and viruses (including non-enveloped viruses)(Omidbakhsh & Sattar, 2006). After control measures were initiated, the outbreak slowed and cases became sporadic, except for at Shelter 3 in January (due to communal housing). Each time control measures failed, the failure was traced to a contaminated environment or fomite.

### PCR pathogen screening

Seventeen fecal samples from cats in Shelters 2 and 3 (collected after the outbreak was recognized) were available for further analysis. Twelve samples representing 15 cats (i.e. including two pools of multiple samples) were subjected to a comprehensive IDEXX feline diarrhea panel pathogen screen (see materials and methods). FECV was detected in one sample, Giardia DNA in 3 samples, and Clostridium perfringens alpha toxin DNA in 6 out of 11 cats that tested, while Clostridium perfringens enterotoxin results were negative for all animals. Clostridium perfringens is part of the normal feline intestinal flora; diarrheic and clinically healthy cats shed the organism at similar rates (Marks, Rankin, Byrne, & Weese, 2011). Type A, the most commonly isolated biotype, produces alpha-toxin and is commonly detected in feces of both diarrheic and healthy dogs (Gizzi et al., 2014). It is not considered a primary cause of diarrhea in cats, but may contribute to diarrhea in cases where a disturbance in the intestinal microenvironment has occurred, such as due to concurrent pathogen infection (Adam, 2014). Giardia is a protozoal parasite that can infect multiple mammalian species and can be associated with diarrhea (and rarely, vomiting) in cats (Janeczko & Griffin, 2010). It has a prepatent period of 5-16 days and while it can cause shelter outbreaks, it would not be capable of causing disease only 24 hours after exposure (Janeczko & Griffin, 2010); a shelter outbreak of Giardia would have a more indolent course. FRV was negative for all samples tested, only one sample (from a pooled litter) was positive for FPV. These tests ruled out the primary bacterial differentials for a shelter outbreak with a short incubation period. None of the positive organisms were considered the main cause of the vomiting and diarrhea outbreak.

### Viral metagenomics analysis

In order to determine the pathogens that caused the outbreak, all 17 available fecal samples were analyzed using viral metagenomics. Viral sequences assigned to five main viral families (*Anelloviridae, Parvoviridae, Papillomaviridae, Polyomaviridae* and *Caliciviridae*) were detected in these fecal samples (Table 3).

**Table 3.**
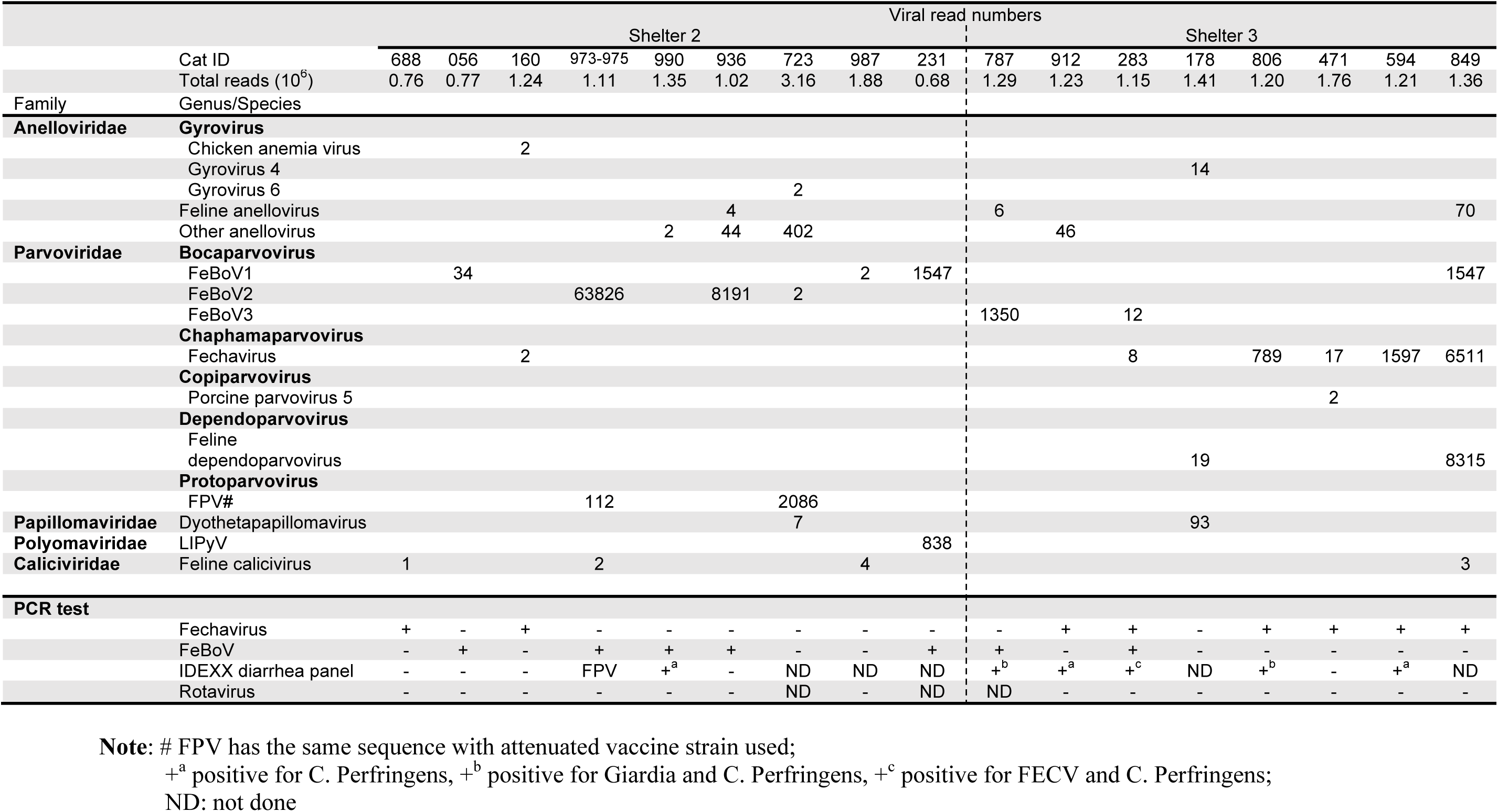
Metagenomic results and PCR test of each case cat.

|  |  | Viral read numbers |  |  |  |  |  |  |  |  |  |  |  |  |  |  |  |  |  |
| --- | --- | --- | --- | --- | --- | --- | --- | --- | --- | --- | --- | --- | --- | --- | --- | --- | --- | --- | --- |
|  |  | Shelter 2 |  |  |  |  |  |  |  |  |  | Shelter 3 |  |  |  |  |  |  |  |
| Cat ID |  | 688 | 056 | 160 | 973-975 | 990 | 936 | 723 | 987 | 231 |  | 787 | 912 | 283 | 178 | 806 | 471 | 594 | 849 |
| Total reads (10<sup>6</sup>) |  | 0.76 | 0.77 | 1.24 | 1.11 | 1.35 | 1.02 | 3.16 | 1.88 | 0.68 |  | 1.29 | 1.23 | 1.15 | 1.41 | 1.20 | 1.76 | 1.21 | 1.36 |
| Family | Genus/Species |  |  |  |  |  |  |  |  |  |  |  |  |  |  |  |  |  |  |
| **Anelloviridae** | **Gyrovirus** |  |  |  |  |  |  |  |  |  |  |  |  |  |  |  |  |  |  |
|  | Chicken anemia virus |  |  | 2 |  |  |  |  |  |  |  |  |  |  |  |  |  |  |  |
|  | Gyrovirus 4 |  |  |  |  |  |  |  |  |  |  |  |  |  | 14 |  |  |  |  |
|  | Gyrovirus 6 |  |  |  |  |  |  | 2 |  |  |  |  |  |  |  |  |  |  |  |
|  | Feline anellovirus |  |  |  |  |  | 4 |  |  |  |  | 6 |  |  |  |  |  |  | 70 |
|  | Other anellovirus |  |  |  |  | 2 | 44 | 402 |  |  |  |  | 46 |  |  |  |  |  |  |
| **Parvoviridae** | **Bocaparvovirus** |  |  |  |  |  |  |  |  |  |  |  |  |  |  |  |  |  |  |
|  | FeBoV1 |  | 34 |  |  |  |  |  | 2 | 1547 |  |  |  |  |  |  |  |  | 1547 |
|  | FeBoV2 |  |  |  | 63826 |  | 8191 | 2 |  |  |  |  |  |  |  |  |  |  |  |
|  | FeBoV3 |  |  |  |  |  |  |  |  |  |  | 1350 |  | 12 |  |  |  |  |  |
|  | **Chaphamaparvovirus** |  |  |  |  |  |  |  |  |  |  |  |  |  |  |  |  |  |  |
|  | Fechavirus |  |  | 2 |  |  |  |  |  |  |  |  |  | 8 |  | 789 | 17 | 1597 | 6511 |
|  | **Copiparvovirus** |  |  |  |  |  |  |  |  |  |  |  |  |  |  |  |  |  |  |
|  | Porcine parvovirus 5 |  |  |  |  |  |  |  |  |  |  |  |  |  |  |  | 2 |  |  |
|  | **Dependoparvovirus** |  |  |  |  |  |  |  |  |  |  |  |  |  |  |  |  |  |  |
|  | Feline dependoparvovirus |  |  |  |  |  |  |  |  |  |  |  |  |  | 19 |  |  |  | 8315 |
|  | **Protoparvovirus** |  |  |  |  |  |  |  |  |  |  |  |  |  |  |  |  |  |  |
|  | FPV# |  |  |  | 112 |  |  | 2086 |  |  |  |  |  |  |  |  |  |  |  |
| **Papillomaviridae** | Dyothetapapillomavirus |  |  |  |  |  |  | 7 |  |  |  |  |  |  | 93 |  |  |  |  |
| **Polyomaviridae** | LIPyV |  |  |  |  |  |  |  |  | 838 |  |  |  |  |  |  |  |  |  |
| **Caliciviridae** | Feline calicivirus | 1 |  |  | 2 |  |  |  | 4 |  |  |  |  |  |  |  |  |  | 3 |
| **PCR test** |  |  |  |  |  |  |  |  |  |  |  |  |  |  |  |  |  |  |  |
|  | Fechavirus | + | - | + | - | - | - | - | - | - |  | - | + | + | - | + | + | + | + |
|  | FeBoV | - | + | - | + | + | + | - | - | + |  | + | - | + | - | - | - | - | - |
|  | IDEXX diarrhea panel | - | - | - | FPV | +<sup>a</sup> | - | ND | ND | ND |  | +<sup>b</sup> | +<sup>a</sup> | +<sup>c</sup> | ND | +<sup>b</sup> | - | +<sup>a</sup> | ND |
|  | Rotavirus | - | - | - | - | - | - | ND | - | ND |  | ND | - | - | - | - | - | - | - |

**Note:** # FPV has the same sequence with attenuated vaccine strain used;
+<sup>a</sup> positive for C. Perfringens, +<sup>b</sup> positive for Giardia and C. Perfringens, +<sup>c</sup> positive for FECV and C. Perfringens;
ND: not done

### Anelloviruses, Papillomaviruses and Polyomavirus

Feline anellovirus were detected in 3 cats, and unclassified anelloviruses were detected in another 4 cats. Anelloviruses are widely prevalent in humans and other mammals, and no clear etiological role of anelloviruses has been identified (Y. Li et al., 2019; Moustafa et al., 2017). Lyon-IARC polyomavirus and feline papillomavirus (Dyothetapappillomavirus 1) were shed by one and two cats, respectively with both genomes showing nucleotide identity of ∼ 99% with those genomes previously reported in cats (Fahsbender, Altan, Estrada, et al., 2019; W. Zhang et al., 2014). Lyon-IARC DNA was previously shown to be shed by 3/5 cat feces in a hoarding situation diarrhea outbreak (one of which was co-infected with FPV) (Fahsbender, Altan, Estrada, et al., 2019; W. Zhang et al., 2014).

### Dietary contamination

Chicken anemia virus (CAV), and gyrovirus 4 and 6 DNA from family *Anelloviridae* were each detected in one cat. CAV is a highly prevalent chicken pathogen (Schat, 2009) and gyroviruses have been frequently detected in human feces presumably originating from consumed infected chicken. (Gia Phan et al., 2013; Niu et al., 2019). Also detected were two sequence reads with a perfect match to porcine parvovirus 5 (*Protoparvovirus* genus) also likely from consumed food (Xiao, Gimenez-Lirola, Jiang, Halbur, & Opriessnig, 2013).

### Vaccine derived sequences

All cats were vaccinated subcutaneously upon entry into the three facilities with Felocell 3 vaccine containing live attenuated FPV, FCV (mainly associated with respiratory systems in cats (Radford, Coyne, Dawson, Porter, & Gaskell, 2007), and feline herpesvirus-1. That vaccine was also sequenced using metagenomics yielding the near complete genome of the FCV (deposited in GenBank MN868063), approximately 85.8% of the FPV genome, and more than 20,000 reads of the herpes virus. Felocell 3 vaccine derived FPV and FCV sequences were compared to those detected by metagenomics in two and four of the diarrheic cats respectively. The sequence reads in the two cats shedding FPV showed 99%-100% identity to the vaccine sequences. The few reads of FCV (1 to 4 sequences reads per sample) also showed a high level of similarity (99-100%) to the sequenced vaccine FCV genome. Given the extensive sequence diversity of FPV and feline FCV (Abd-Eldaim, Potgieter, & Kennedy, 2005; Balboni et al., 2018; Battilani et al., 2011; Hou et al., 2016) we conclude that the FPV and FCV sequence reads detected in feces by metagenomics were vaccine derived.

### Parvoviruses

Viral sequences belonging to the *Bocaparvovirus*, and *Chaphamaparvovirus* genera were the most abundant and detected in 14/17 cats (Table 3). Three different feline bocaparvoviruses (FeBoV1-3) were shed by 3, 3 and 2 cats respectively. We then used a pan-FeBoV PCR to test these 17 samples. One cat, FeBoV negative by metagenomics, was PCR positive (FeBoV2 by Sanger sequencing of PCR amplicon). Two other cats positive for bocaviruses with only 2 sequence reads were negative by PCR. All other PCR results were consistent with metagenomics results (Table 3). All together FeBoV DNA was detected by metagenomics and/or PCR in 9/17 animals.

Six cats also yielded sequence reads related to multiple parvoviruses in the *Chaphamaparvovirus* genus. Using de novo assembly and specific PCR to fill gaps, two near full-length genomes of 4225 and 4134 bases (Figure 3A) were generated from cats #594 and #849 (GenBank accession numbers MN396757 and MN794869). The two genomes share 99.1% identity of NS1 and 99.6% identity of VP1 at protein level, and 99.2% and 99.3% at nucleotide level. These genomes encoded the two main ORFs shared by all parvoviruses, a 658aa ORF encoding non-structural replication protein (NS1), and a 508aa ORF encoding viral capsid (VP1). The closest relative to these genomes was a parvovirus recently reported in canine feces named cachavirus (MK448316)(Fahsbender, Altan, Seguin, et al., 2019), with a nucleotide identity of 74.2% and NS1 and VP1 protein identity of 65.5% and 68.8% respectively. Other major ORFs consisted of a 186aa ORF within the NS1 labeled NP and a potential 248aa ORF overlapping the 5’-UTR and NS1. The NP ORF is widely conserved in chaphamaparvoviruses (Christopher J. Jolly, 2019; Roediger et al., 2018). No significant similarity was found for the 248aa ORF.

**Figure 3.**
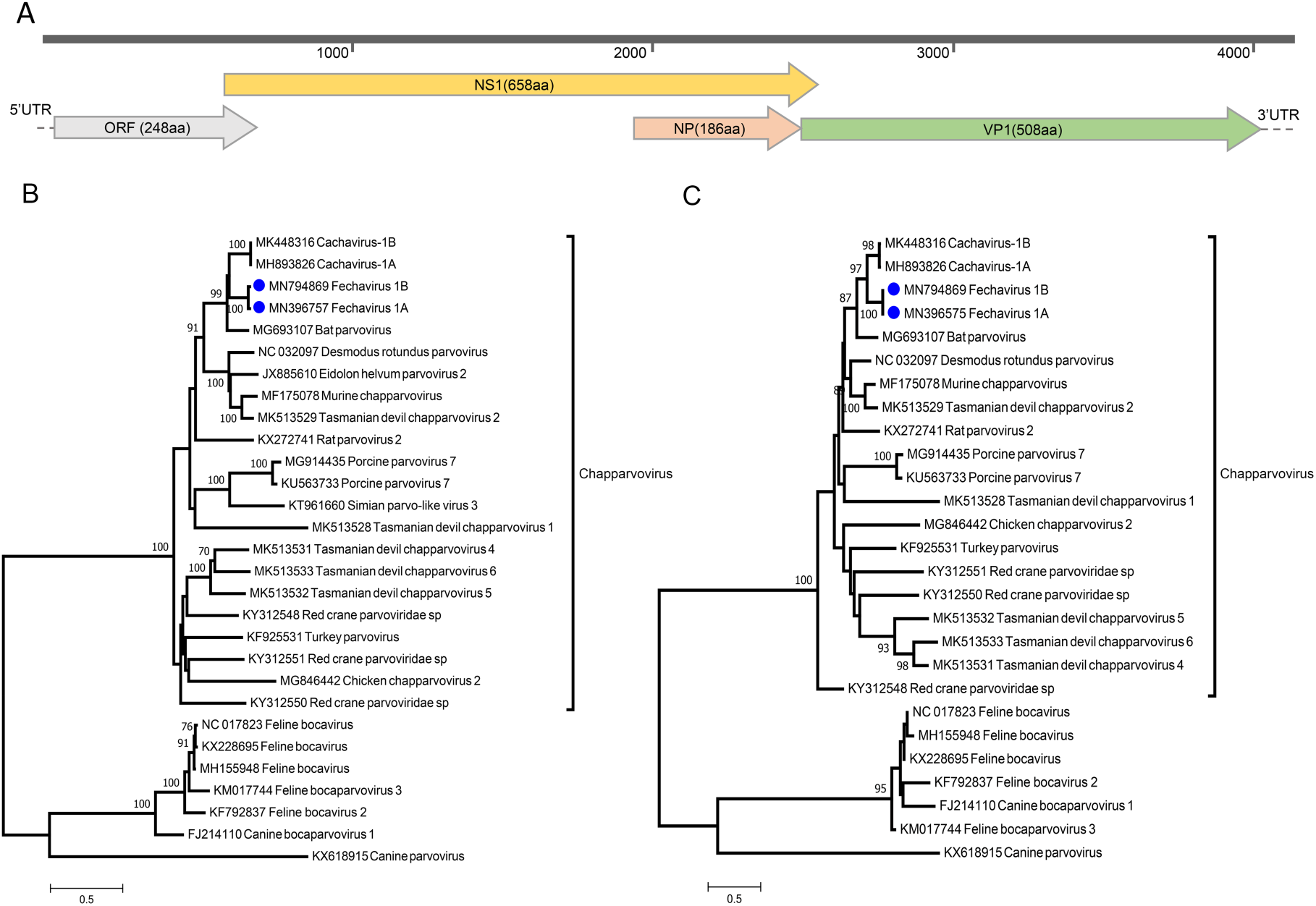
A) Genome organization of the fechavirus. Phylogenetic tree of NS1 protein sequences (B) and VP1 protein sequences (C) of chapparvoviruses. Scale bar, 0.5 amino acid substitutions per site.

Under an April 2019 ICTV proposal updating the latest published classification (Cotmore et al., 2019) members of the same parvovirus species should show >85% identity for NS1. The fechavirus genome therefore qualifies as a member of a new species in the proposed *Chaphamaparvovirus* genus. We named this novel virus <u>fe</u>line <u>cha</u>phamaparvo<u>virus</u> or fechavirus. Phylogenetic analysis confirmed the closest relative to be the canine cachavirus in both NS and VP ORFs (Figure 3B and C).

Specific PCR primers were designed based on the fechavirus genomes and used to test all 17 fecal samples. Besides the six fechavirus positive fecal samples detected by NGS, two more samples were positive by PCR. All four vomit samples tested (from unknown cats in shelter 3) were also PCR positive. A total of 8 animals therefore shed fechavirus of which one was co-infected with FeBoV3 (Cat #283). Of the 17 animals available for analyzes only one was negative for both FeBoV and fechavirus DNA (cat # 178).

Longitudinally collected fecal samples were collected from 4 cats and PCR tested for fechavirus DNA. Cat #688 was fechavirus positive on the last day of his illness; while cats #912 and #283 were positive for a single time point while disease signs were manifest for a further 4 or 9 more days. Cat #594 shed fechavirus over 4 days while symptomatic plus a further 7 days after recovery (Table 4).

**Table 4.** Serial fechavirus results from cats tested on multiple dates.

| ID | D1 | D2 | D3 | D4 | D5 | D6 | D7 | D8 | D9 | D10 | D11 | D12 |
| --- | --- | --- | --- | --- | --- | --- | --- | --- | --- | --- | --- | --- |
| 688 | + |  | - |  |  |  |  |  |  |  |  |  |
| 912 | + |  | - |  |  | - | - |  |  |  |  |  |
| 594 | + |  | + | + | + | + | + |  |  | + |  | + |
| 283 | - | + | - |  |  |  |  |  |  |  |  |  |

Note: Days (D1-D12) reflect day samples were collected. Days of illness are shaded starting at first sample collection. “+” and “-” means PCR positive or negative for the fechavirus. Illness onset was 10, 2, 2, 3 days before Day1 for cats #688, 912, 594 and 283, respectively.

Dependoparvovirus sequence reads were also found in two cats. A near full-length genome of 4315 bases could be assembled from cat #849 (also infected with fechavirus) whose phylogenetic analysis showed it to be related to a recently reported dependoparvovirus genome from bats (*Desmodus rotundus*) sharing NS1 and VP1 protein with 51.6% and 46.7% identities (Supplementary Figure 1).

### Epidemiology

The timeline of disease presentation and fecal shedding status of key animals in the transmission chain between the 3 facilities was determined. There were no samples from cats in Shelter 1 available for testing, but cat #688 was fechavirus positive shortly after being housed in the same area as cat #853 and #614, who were sick upon transfer from Shelter 1 to Shelter 2. A second cat in shelter 2 was also shedding fechavirus (cat #160). Of the other 15 affected cats in Shelter 2 a total of 5 individual cats and a sample pool (mixed feces from 3 cats) were FeBoV1, 2, or 3 positive. Fechavirus positive cat #160 was then in direct contact with subsequently diseased cat #992 from shelter 3. Despite clean breaks imposed by cleaning and emptying the facility in shelter 3 the disease continued to recur and spread. Of the seven cat fecal samples available for study from shelter 3 fechavirus was detected in five and bocaparvovirus in 2 animals. Only a single case yielded both fechavirus and bocaparvovirus DNA (#283). A further four vomit samples from unknown cat(s) in shelter 3 were all fechavirus PCR positive. Last, an initially healthy cat in Shelter 2 developed disease signs immediately after cats from Shelter 3 were returned to Shelter 2 during the last wave of the outbreak and was found to be fechavirus positive (cat #849).

## Discussion

Currently identified feline parvoviruses belong to two genera of the *Parvoviridae* family, namely three bocaparvoviruses species, FeBoV1-3 (Takano, Takadate, Doki, & Hohdatsu, 2016; Yi et al., 2018; W. Zhang et al., 2014), and two protoparvovirus species, feline panleukopenia virus (FPV)(Barrs, 2019) and feline bufavirus (FBuV) (Diakoudi et al., 2019). FeBoV1 was first discovered in in multiple tissues of cats in Hong Kong (Lau et al., 2012) and subsequently reported in cat feces in the US, Japan, Europe and China (Ng et al., 2014; Takano et al., 2016; Yi et al., 2018; W. Zhang et al., 2014). Similarly to human bocaparvovirus DNA commonly found in the feces of healthy humans (Jartti et al., 2012; Kapoor et al., 2010; Ong, Schuurman, & Heikens, 2016), frequent feline bocaparvovirus DNA detection in healthy cats raise questions regarding its pathogenicity in cats (Diakoudi et al., 2019; Lau et al., 2012; Ng et al., 2014; Piewbang, Kasantikul, Pringproa, & Techangamsuwan, 2019; W. Zhang et al., 2014). A recent study showed a possible association between FBoV-1 infection, potentially aggravated by FPV, and hemorrhagic enteritis (Diakoudi et al., 2019; Lau et al., 2012; Ng et al., 2014; Piewbang et al., 2019; W. Zhang et al., 2014). FPV is an extensively studied pathogen that can lead to reduction in circulating white blood cell and enteritis (Barrs, 2019; Truyen & Parrish, 2013). The pathogenicity of FeBuV in cats is currently unknown (Diakoudi et al., 2019; Stuetzer & Hartmann, 2014) but the virus shares very high sequence identity with a canine bufavirus (Diakoudi et al., 2019; J. Li et al., 2019; Martella et al., 2018).

Using metagenomics, we found FeBoV1, 2, and 3 and a novel chaphamaparvovirus we named fechavirus in a large fraction of fecal samples and fechavirus in all vomit samples from sick cats in a multi-facility outbreak. Subsequent PCR testing confirmed the presence of either a bocavirus or fechavirus or both (n=1 cat #283) of these viruses in all but one of the seventeen sick cats tested (cat #178). The outbreak in shelter 2 predominantly tested positive for FeBoV (7/9 FeBoV +ve and 2/9 fechavirus +ve) while the sick cats in shelter 3 were mainly shedding fechavirus (6/7 fechavirus +ve, 2/7 FeBov +ve). Another cat housed in shelter 2 who developed vomiting after sick cats were transferred back from Shelter 3 was also shedding fechavirus as well as a novel dependovirus. The co-detection of these viral genomes indicate that fechavirus may provide the required help for replication-defective dependoviruses.

FPV and feline FCV reads were determined to originate from recently inoculated attenuated vaccine strains while other viruses were deemed to be asymptomatic infections, derived from chicken and pork viruses in consumed food, or detected only in sporadic cases. FCV sequences were negative by PCR, this could be the results of low viral load in those cats.

The close genetic relationship of fechavirus to cachavirus found in both diarrheic and healthy dog feces (Fahsbender, Altan, Seguin, et al., 2019), may reflects a cross-carnivore virus transmission as occurred with FPV mutating into the highly pathogenic canine parvovirus (Allison et al., 2013; Hoelzer, Shackelton, Parrish, & Holmes, 2008).

Daily collected fecal samples from symptomatic cats revealed transient shedding for 3/4 animals with fechavirus DNA only detected at a single time point. One animal shed fechavirus DNA during all twelve days of sampling both while exhibiting disease signs and seven days after disease resolution. The short duration of fechavirus shedding for 3/4 animals, also typical of FPV shedding (Bergmann et al., 2019), may account for its non-detection in some of the affected animals in shelter 2 and 3. All but one of the fechavirus negative samples were positive for one of three different feline bocavirus indicating that both bocaviruses and fechavirus were circulating in these shelters. Because of the high diversity of bocaviruses found here, belonging to three distinct species, and the typically asymptomatic nature of FeBoV infections, a role for bocaviruses in the multi-facility transmission of this disease outbreak seems unlikely. A pathogenic role for fechavirus seems more likely as viral DNA was detected in key animals with contacts across the three facilities experiencing outbreaks with similar disease signs.

A key clinical attribute of this unusual outbreak was the prominence of indirect routes of transmission, including ongoing fomite transmission after robust control measures were implemented. Another notable characteristic of this case was the very short minimum incubation period. Combined with the absence of any known pathogen identified on routine screening, this points to a novel non-enveloped enteric virus as the most likely causative agent. Animal shelters and veterinary hospitals should maintain good routine infection control procedures and consider novel viruses, including fechavirus, in the diagnosis of feline gastrointestinal disease.

In this study, we characterized the virome from a cat diarrhea and vomiting outbreak, and showed that besides different FeBoV, a new chaphamaparvovirus was associated with gastrointestinal disease. Future studies on its prevalence, genetic diversity, and tissue distribution in healthy and diarrheic cats are needed to further investigate a possible etiologic role in feline diseases.

## Author Contributions

Conceptualization, YL, EG, ED; Data curation, YL, EG, AI, EA, MAS, ME, XD; Formal analysis, YL, EG, ED; Methodology, YL, EG; Supervision, ED, EG; Writing original draft, YL, EG, ED; Writing review and editing, YL, EG, ED.

## Acknowledgments

The authors would like to acknowledge Melissa Beall, Roxanne Chan, and Phyllis Tyrrel from IDEXX for helpful comments and assistance with clinical data. Funding from IDEXX and the Vitalant Research Institute.

## Ethics Statement

The authors confirm that the ethical policies of the journal, as noted on the journal’s author guidelines, have been adhered to. No ethical approval was required as samples were collected as part of routine outbreak investigation.

## Data Availability Statement

The data that supports the findings of this study are available in the main body of this article.

## Figures

**Supplemental table 1:**
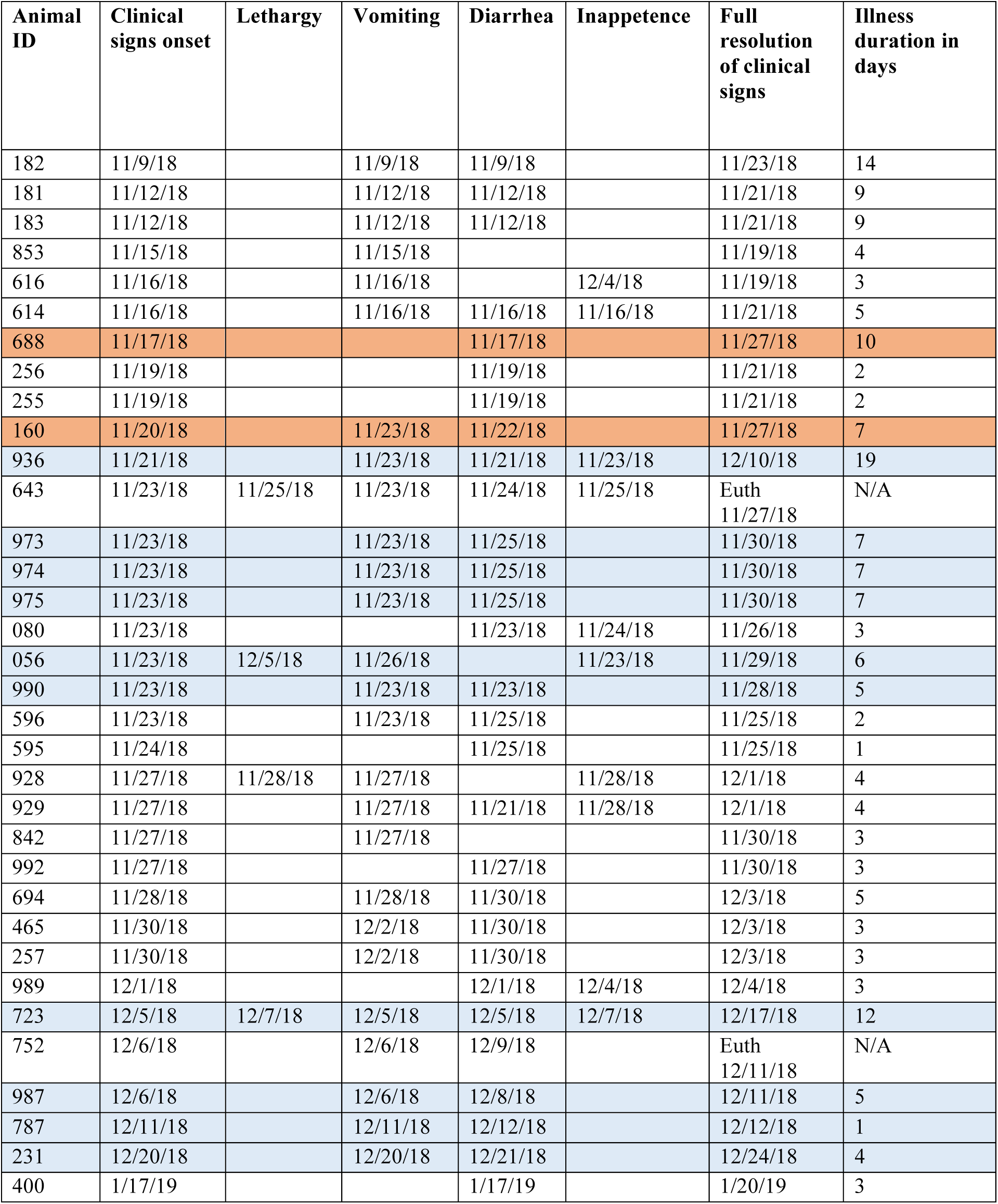
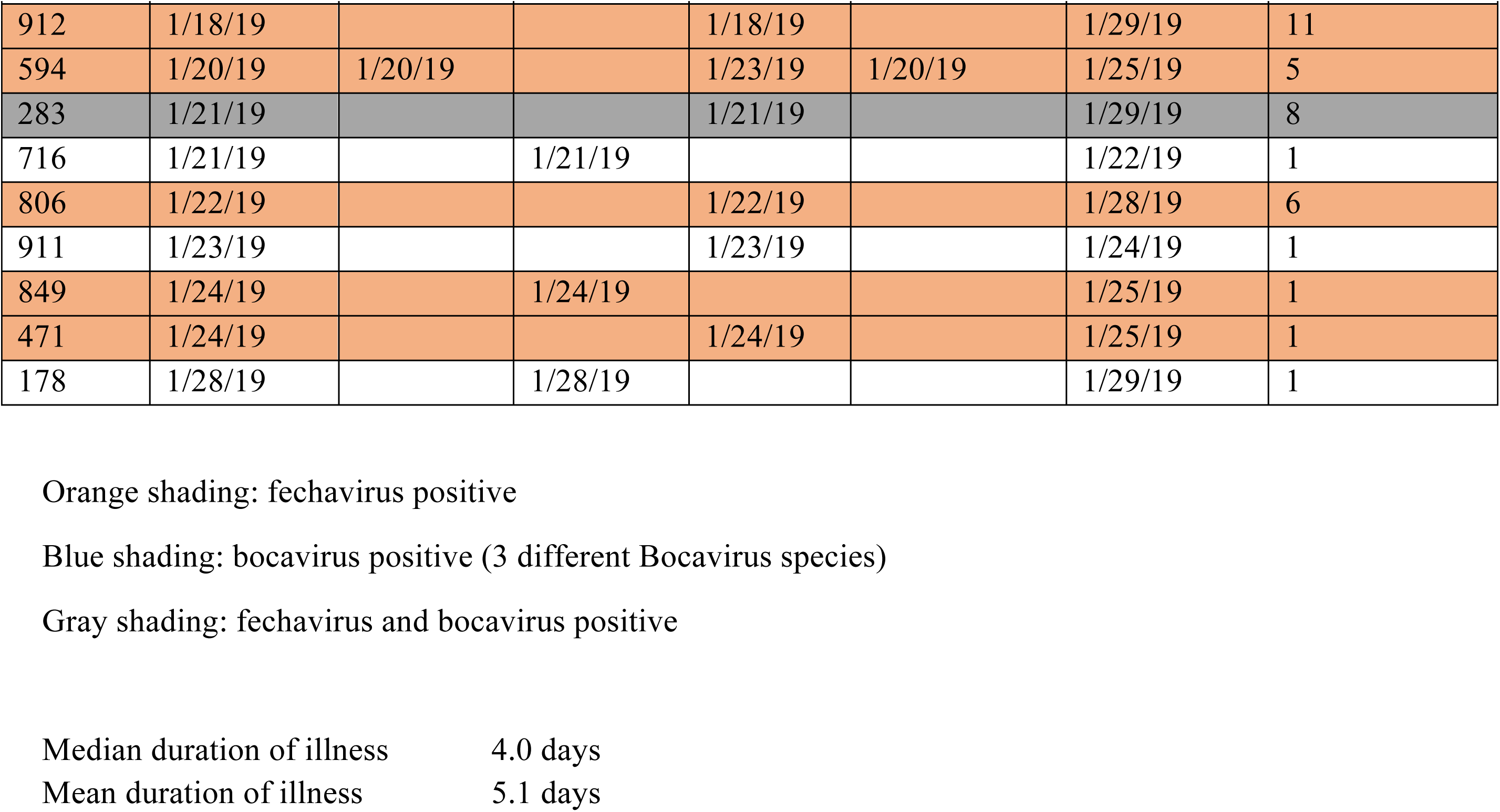
Clinical signs and case definition ratings for all cats meeting case definition

| **Animal ID** | **Clinical signs onset** | **Lethargy** | **Vomiting** | **Diarrhea** | **Inappetence** | **Full resolution of clinical signs** | **Illness duration in days** |
| --- | --- | --- | --- | --- | --- | --- | --- |
| 182 | 11/9/18 |  | 11/9/18 | 11/9/18 |  | 11/23/18 | 14 |
| 181 | 11/12/18 |  | 11/12/18 | 11/12/18 |  | 11/21/18 | 9 |
| 183 | 11/12/18 |  | 11/12/18 | 11/12/18 |  | 11/21/18 | 9 |
| 853 | 11/15/18 |  | 11/15/18 |  |  | 11/19/18 | 4 |
| 616 | 11/16/18 |  | 11/16/18 |  | 12/4/18 | 11/19/18 | 3 |
| 614 | 11/16/18 |  | 11/16/18 | 11/16/18 | 11/16/18 | 11/21/18 | 5 |
| 688 | 11/17/18 |  |  | 11/17/18 |  | 11/27/18 | 10 |
| 256 | 11/19/18 |  |  | 11/19/18 |  | 11/21/18 | 2 |
| 255 | 11/19/18 |  |  | 11/19/18 |  | 11/21/18 | 2 |
| 160 | 11/20/18 |  | 11/23/18 | 11/22/18 |  | 11/27/18 | 7 |
| 936 | 11/21/18 |  | 11/23/18 | 11/21/18 | 11/23/18 | 12/10/18 | 19 |
| 643 | 11/23/18 | 11/25/18 | 11/23/18 | 11/24/18 | 11/25/18 | Euth<br>11/27/18 | N/A |
| 973 | 11/23/18 |  | 11/23/18 | 11/25/18 |  | 11/30/18 | 7 |
| 974 | 11/23/18 |  | 11/23/18 | 11/25/18 |  | 11/30/18 | 7 |
| 975 | 11/23/18 |  | 11/23/18 | 11/25/18 |  | 11/30/18 | 7 |
| 080 | 11/23/18 |  |  | 11/23/18 | 11/24/18 | 11/26/18 | 3 |
| 056 | 11/23/18 | 12/5/18 | 11/26/18 |  | 11/23/18 | 11/29/18 | 6 |
| 990 | 11/23/18 |  | 11/23/18 | 11/23/18 |  | 11/28/18 | 5 |
| 596 | 11/23/18 |  | 11/23/18 | 11/25/18 |  | 11/25/18 | 2 |
| 595 | 11/24/18 |  |  | 11/25/18 |  | 11/25/18 | 1 |
| 928 | 11/27/18 | 11/28/18 | 11/27/18 |  | 11/28/18 | 12/1/18 | 4 |
| 929 | 11/27/18 |  | 11/27/18 | 11/21/18 | 11/28/18 | 12/1/18 | 4 |
| 842 | 11/27/18 |  | 11/27/18 |  |  | 11/30/18 | 3 |
| 992 | 11/27/18 |  |  | 11/27/18 |  | 11/30/18 | 3 |
| 694 | 11/28/18 |  | 11/28/18 | 11/30/18 |  | 12/3/18 | 5 |
| 465 | 11/30/18 |  | 12/2/18 | 11/30/18 |  | 12/3/18 | 3 |
| 257 | 11/30/18 |  | 12/2/18 | 11/30/18 |  | 12/3/18 | 3 |
| 989 | 12/1/18 |  |  | 12/1/18 | 12/4/18 | 12/4/18 | 3 |
| 723 | 12/5/18 | 12/7/18 | 12/5/18 | 12/5/18 | 12/7/18 | 12/17/18 | 12 |
| 752 | 12/6/18 |  | 12/6/18 | 12/9/18 |  | Euth<br>12/11/18 | N/A |
| 987 | 12/6/18 |  | 12/6/18 | 12/8/18 |  | 12/11/18 | 5 |
| 787 | 12/11/18 |  | 12/11/18 | 12/12/18 |  | 12/12/18 | 1 |
| 231 | 12/20/18 |  | 12/20/18 | 12/21/18 |  | 12/24/18 | 4 |
| 400 | 1/17/19 |  |  | 1/17/19 |  | 1/20/19 | 3 |
| 912 | 1/18/19 |  |  | 1/18/19 |  | 1/29/19 | 11 |
| 594 | 1/20/19 | 1/20/19 |  | 1/23/19 | 1/20/19 | 1/25/19 | 5 |
| 283 | 1/21/19 |  |  | 1/21/19 |  | 1/29/19 | 8 |
| 716 | 1/21/19 |  | 1/21/19 |  |  | 1/22/19 | 1 |
| 806 | 1/22/19 |  |  | 1/22/19 |  | 1/28/19 | 6 |
| 911 | 1/23/19 |  |  | 1/23/19 |  | 1/24/19 | 1 |
| 849 | 1/24/19 |  | 1/24/19 |  |  | 1/25/19 | 1 |
| 471 | 1/24/19 |  |  | 1/24/19 |  | 1/25/19 | 1 |
| 178 | 1/28/19 |  | 1/28/19 |  |  | 1/29/19 | 1 |

Orange shading: fechavirus positive
Blue shading: bocavirus positive (3 different Bocavirus species)
Gray shading: fechavirus and bocavirus positive
Median duration of illness 4.0 days
Mean duration of illness 5.1 days

**Figure S1.**
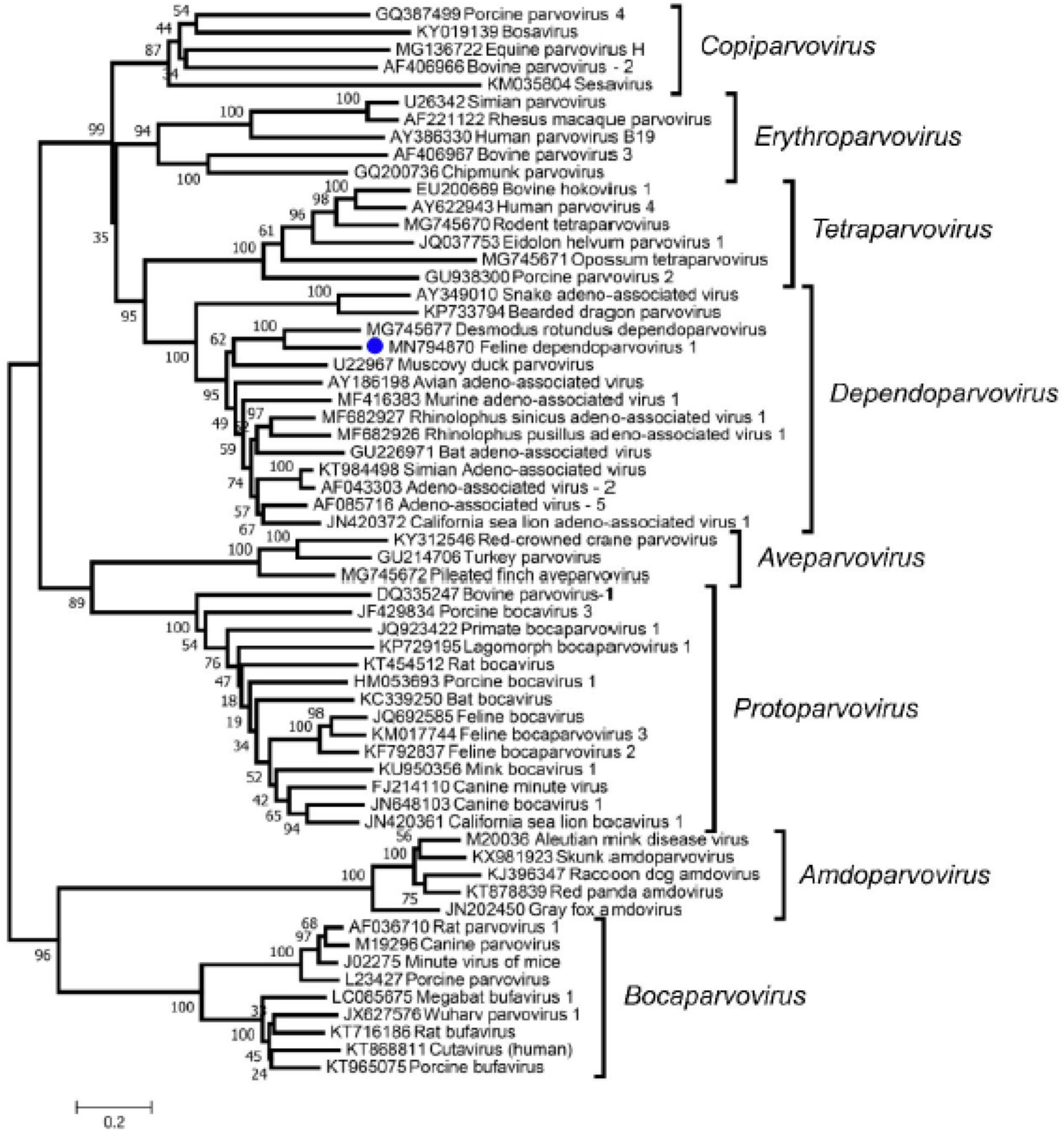
Phylogenetic tree of the complete NS1 protein sequences of subfamily *Parvovirinae*. The new dependoparvovirus identified in cat is shown with a blue circle.

